# Evaluating fMRI-Based Estimation of Eye Gaze during Naturalistic Viewing

**DOI:** 10.1101/347765

**Authors:** Jake Son, Lei Ai, Ryan Lim, Ting Xu, Stanley Colcombe, Alexandre Rosa Franco, Jessica Cloud, Stephen LaConte, Jonathan Lisinski, Arno Klein, R. Cameron Craddock, Michael Milham

## Abstract

The collection of eye gaze information during functional magnetic resonance imaging (fMRI) is important for monitoring variations in attention and task compliance, particularly for naturalistic viewing paradigms (e.g., movies). However, the complexity and setup requirements of current in-scanner eye-tracking solutions can preclude many researchers from accessing such information. Predictive eye estimation regression (PEER) is a previously developed support vector regression-based method for retrospectively estimating eye gaze from the fMRI signal in the eye’s orbit using a 1.5-minute calibration scan. Here, we provide confirmatory validation of the PEER method’s ability to infer eye gaze on a TR-by-TR basis during movie viewing, using simultaneously acquired eye tracking data in five individuals (median angular deviation < 2°). Then, we examine variations in the predictive validity of PEER models across individuals in a subset of data (n=448) from the Child Mind Institute Healthy Brain Network Biobank, identifying head motion as a primary determinant. Finally, we accurately classify which of two movies is being watched based on the predicted eye gaze patterns (area under the curve = .90 ± .02) and map the neural correlates of eye movements derived from PEER. PEER is a freely available and easy-to-use tool for determining eye fixations during naturalistic viewing.

## INTRODUCTION

A classic challenge for functional magnetic resonance imaging (fMRI) is the identification of variations in attention, arousal and compliance with task demands during a given scan (Church, Petersen, and Schlaggar 2010; Price and Friston 1999; Brown et al. 2005; K. Murphy and Garavan 2004). This is particularly true when participants are required to perform tasks that are tedious or tiresome. In studies requiring active task performance during fMRI scans, these concerns can be addressed in part by analyzing performance measures (e.g., accuracy, reaction time) (Koten et al. 2013). However, such solutions do not work for studies of passive states (e.g., resting state fMRI, naturalistic viewing fMRI), as there are no such responses to monitor. The resting state fMRI literature has struggled with this issue due to the increased likelihood of individuals falling asleep, including micro-naps, (Tagliazucchi and Laufs 2014; J. Wang et al. 2017; Laumann et al. 2017), driving most to require participants to remain with their eyes open during scans - a requirement that can be tracked through direct observation by a technician or via video recording. The requirements of conducting fMRI movie viewing experiments are greater, as one needs to know that an individual is actually paying attention to specific information on the screen.

Eye tracking devices provide an obvious solution to the additional challenges inherent to naturalistic viewing fMRI, as they can provide high fidelity fixation information from which the level of participant engagement and preferential viewing patterns can be readily inferred (Hansen and Ji 2010; K. Murphy and Garavan 2004). A growing number of systems are available for use in the MRI environment, meeting ferromagnetic and radiofrequency requirements and overcoming logistical requirements regarding their positioning (e.g., long-range mounts) (Kimmig et al. 1999; Kanowski et al. 2007; Ferri et al. 2013); costs for these systems are slowly becoming less prohibitive. However, these devices remain outside the range of access for many due to the added layers of complexity (e.g. operator training, setup time, synchronization and analysis of an additional data type) that can be burdensome.

An alternative solution is predictive eye estimation regression (PEER) (LaConte et al. 2006, 2007), an imaging-based method that uses machine learning algorithms (i.e., support vector regression [SVR] (Drucker et al. 1996)) to estimate the direction of gaze during each repetition (TR) in the fMRI time series based on voxel-wise data from the eyes. The method leverages variations in the MRI contrast within the eye to make its predictions, likely between the vitreous and aqueous humors (Fanea and Fagan 2012). Prior work has validated this approach by comparing PEER-derived fixations with simultaneously acquired eye tracking (LaConte et al. 2007), as well as with perimetry (Adelore 2008). However, PEER has only been used in a few studies to date, which were small in size and tended to focus on neurologically normal adults (LaConte et al. 2006; Sathian et al. 2011; LaConte et al. 2007; Adelore 2008).

Here, we provide a multifaceted assessment of the readiness of the PEER technique for application in movie viewing fMRI studies. First, we provide a confirmatory validation of prior reports regarding the PEER method, using simultaneous eye tracking and fMRI data obtained during movie viewing in five adults at the Nathan Kline Institute (NKI) to establish concurrent validity. Then, we used the large-scale Child Mind Institute Healthy Brain Network Biobank (HBN) (Alexander et al. 2017) to examine factors that can impact the predictive validity of the PEER models generated for individuals, using a subset of fMRI data (n=448). The availability of 2-3 PEER scans for each participant allowed us to examine variations in the predictive validity and reproducibility of PEER models, as well as minimum data requirements. The inclusion of children of varying ages (5-21 years old) and head motion during scan sessions allowed us to assess potential sources of artifacts related to compliance (Kotsoni, Byrd, and Casey 2006; Wilke et al. 2003; Schlund et al. 2011; Alexander et al. 2017; Gaillard, Grandin, and Xu 2001; Satterthwaite et al. 2012).

In addition to PEER calibration scans, two movie fMRI scans were available for most participants of the HBN sample (*Despicable Me, The Present*). These allowed us to characterize the consistency of eye gaze patterns calculated using PEER across participants for a given movie. We also examined the specificity of eye tracking patterns obtained for differing movie stimuli using the predicted eye gaze patterns to train and test a support vector machine classifier. Finally, given that out-of-scanner eye tracking device data were available for *The Present*, we assessed comparability of the PEER-generated fixations to gold standard results.

## MATERIALS AND METHODS

### Participants: Simultaneous fMRI-Eye Tracking Validation Dataset

Five adult participants were recruited at the Nathan Kline Institute using a local advertisement (ages: 27-60; 5M). Participants were screened for MRI contraindications and informed consent was obtained from participants according to institutional guidelines prior to participating in the study.

### Participants: CMI Healthy Brain Network (HBN)

The Healthy Brain Network is a large-scale data collection effort focused on the generation of an open resource for studying pediatric mental health and learning disorders (Alexander et al. 2017). The ongoing data collection contains a range of phenotypes, spanning typical and atypical brain development. We included data from 480 participants (ages: 5.0-21.0; 10.3 ± 3.5) collected at the Rutgers University site, which had the largest number of complete imaging datasets available at the time of our analyses. As outlined in greater detail in the data descriptor publication for the Healthy Brain Network (Alexander et al. 2017), approximately 80 percent of the participants in the sample have one or more diagnosable disorders according to the DSM-5. This dataset includes a high proportion of participants with Attention Deficit Hyperactivity Disorder (ADHD; ∼50%) and children as young as age 5.0. Both of these participant types have a higher likelihood of head motion, allowing us to study its impact on PEER analyses (Kotsoni, Byrd, and Casey 2006; Wilke et al. 2003; Schlund et al. 2011; Alexander et al. 2017).

### Scanners

All data in the present work were obtained using Siemens 3.0T Tim Trio MRI scanners. The simultaneous fMRI-eye tracking data were collected using a Trio scanner based at the Nathan Kline Institute, which was equipped with an in-scanner commercial eye tracking device (EyeLink 1000+; sampling rate = 250Hz). The Healthy Brain Network data used were from the subset collected at the Rutgers University Brain Imaging Center (RUBIC). The same sequence based on the NIH ABCD Study (Casey et al. 2018) was used for all fMRI scans at the two sites (multiband factor 6; 2.4mm isotropic voxel size; TR = 800ms; TE = 30ms).

### Imaging Protocol: Simultaneous fMRI-Eye Tracking Validation Dataset

We obtained simultaneous fMRI and eye tracking data from five adult participants at the Nathan Kline Institute. Data were collected from each participant during two PEER calibration scans (described in detail in the following sections) and a movie viewing scan for *The Present* [0:00-4:19□min; https://vimeo.com/152985022]. The display size was 44.5cm × 34.5 cm, placed at a distance of 134.5cm from the participants’ eyes, and subtending approximately 19 × 14.7 degrees of visual angle.

### Imaging Protocol: CMI Healthy Brain Network

During the imaging session, each participant completed a minimum of two PEER calibration scans (3 scans for n = 430, 2 scans for n = 50). Two movie viewing scans were included as well: *Despicable Me* [10□min clip, DVD version exact times 1:02:09 – 1:12:09] and *The Present* [0:00-3:21□min; HBN participants were not shown the credits]. The display totaled 1680 × 1050 pixels (34.7cm × 21cm). In the scanner, the participants’ eyes were approximately 10 cm from the mirror and the mirror was located 125cm from the screen, yielding a visual angle of 14.2° × 8.9°.

### Predictive Eye Estimation Regression (PEER)

#### PEER Scan Instructions

Participants were asked to fixate on a series of white dots that were sequentially presented, each at one of 25 unique positions (duration = 4 seconds each [5 fMRI repetitions]). A white dot appeared three separate times at the center of the screen, resulting in 27 total fixation points displayed for 4 seconds each, or a time series of 135 points. The positions were selected to ensure coverage of all corners of the screen as well as the center (see Supplementary Figure 1). The PEER calibration scans were distributed throughout the imaging session such that they flank the other scan types (e.g. rest, movie viewing) and allow for a sampling of possible changes in image properties over time.

#### Image Processing

Consistent with prior work (LaConte et al. 2006, 2007), a minimal image processing strategy was employed for the PEER scans. Using the Configurable Pipeline for the Analysis of Connectomes (C-PAC) (Craddock et al. 2013), we performed the following steps: motion correction, image intensity normalization, temporal high-pass filtering (cutoff = 100s), and spatial filtering (FWHM = 6mm). The preprocessed functional data for each participant was then registered to the corresponding high-resolution anatomical T1 image using boundary-based registration via FLIRT (Mark Jenkinson et al. 2002; M. Jenkinson and Smith 2001). The final fMRI data were registered to the MNI152 (Grabner et al. 2006) template space using ANTs (Avants et al. 2011).

#### Quality Assurance

Two researchers visually inspected the middle volume of each participant’s PEER calibration scans for incomplete coverage of the orbit (i.e. missing eye signal), leading to the exclusion of 32 participants from the analysis. At least two PEER scans were available for each participant (3 scans for n = 409, 2 scans for n = 39). Given that *The Present* was added to the HBN imaging protocol later than *Despicable Me*, fewer participants had both scans. We inspected the movie scans and identified 427 *Despicable Me scans* and 360 scans of *The Present* with complete coverage of eye signal. In the following experiments, we removed participants with low quality scans relevant to a given analysis.

#### Eye Mask

We limited each fMRI scan to the region corresponding to the MNI152 eye-mask template. Isolation of signal to the orbit was done for two reasons. First, prior work (LaConte et al. 2006, 2007) suggests that signal from the eyes provides adequate information to predict eye fixations. Second, this would reduce the number of features of the dataset to accelerate model generation and eye gaze estimation using PEER.

#### Averaging Prior to Training

To minimize undesired variations in the training data for a given participant, at each voxel, we: 1) mean-centered and variance-normalized (i.e., z-scored) the time series, and then 2) averaged the five consecutive time points associated with each calibration target presented (reducing the fMRI time series used for training from 135 to 27 points). Our pilot testing for the present work suggested that this latter step improves model performance, likely due to its ability to mitigate the impact of fluctuations in fixation stability that can arise over time (Ditchburn and Ginsborg 1953; Vingolo, Fragiotta, and Cutini 2014; Morales et al. 2016). This averaging process was limited to PEER model training; all applications of the PEER model in the present work were based on individual time point data (i.e., generating a unique prediction for every TR).

#### Model Generation

For each participant, two separate support vector regression models were trained using Scikit Learn (https://scikit-learn.org/stable/) - one for x-direction fixation locations and one for y-direction fixation locations. In accord with prior work (LaConte et al. 2006, 2007), PEER was used to predict the 25 positions using the voxel-wise time series data (i.e., a unique predictor was included for each voxel, with the following parameters: C=100, epsilon=0.01). C represents the regularization parameter that controls the trade-off between errors in the training set and margin maximization. Regression errors smaller than the epsilon threshold (margin of tolerance) are ignored in the calculation of the loss function. The full PEER model code is available here: https://github.com/ChildMindInstitute/pypeer.

#### Assessing Model Validity: Simultaneous fMRI-Eye Tracking Validation Dataset

For each participant, we: 1) trained a PEER-generated SVR model using the first calibration scan (Scan1) and predicted eye gaze from their second calibration scan (Scan2) on a TR-by-TR basis, and 2) segmented the raw data from the in-scanner eye-tracker (sampling rate = 250Hz) to 800 ms windows, to match the sampling rate (TR = 800ms) of the MRI scanner; for each window, the median of the raw samples was used as an estimate of the average direction of eye gaze detected by the eye tracker. PEER model validity was then assessed using Pearson’s correlation coefficient between the model-generated fixation time series and the downsampled eye tracking data, for both the second PEER calibration scan and *The Present*. We also assessed validity by quantifying differences in the degree of visual angle measured by PEER and the in-scanner tracker using the absolute angular deviation (this measure eliminates the positive or negative direction of the difference in gaze).

#### Estimating Model Predictive Validity: CMI Healthy Brain Network

For each participant, the PEER-generated SVR model trained using Scan1 (PEER1) was used to predict eye fixations from their second calibration scan (Scan2). PEER1 model predictive validity was assessed using Pearson’s correlation coefficient between the model-generated fixation time series and the stimulus locations from the calibration sequence. The angular deviation was also calculated for each time point between the model-generated fixation time series and the stimulus locations from the calibration sequence. Pediatric neuroimaging studies demonstrate a decline in scan quality over the course of an imaging session, which implies that the PEER calibration scans (and the resulting SVR models) may differ in overall quality based on the calibration scan’s timing in the imaging session (Kotsoni, Byrd, and Casey 2006; Wilke et al. 2003; Schlund et al. 2011; Kohavi 1995; Refaeilzadeh, Tang, and Liu 2016; Bradley 1997). Thus, we compared the PEER models trained using Scan1, Scan3, or both scans in their ability to predict eye fixations from Scan2. To quantify differences in PEER scan quality, we conducted a paired t-test to compare Scan1 and Scan3 with respect to head motion (i.e. framewise displacement, DVARS) (Power et al. 2014, 2012).

### Factors Associated with PEER Prediction Quality

#### Head Motion and Compliance

Head motion is one of the most consistent sources of artifacts in fMRI analyses (Power et al. 2014, 2012; Satterthwaite et al. 2012). We examined the impact of head motion on PEER model predictive validity using mean framewise displacement and standardized DVARS (Power et al. 2012; Nichols 2017). To do so, we assessed the relationship between measures of motion and the predictive validity of PEER1 (ability to predict fixations from Scan2) via linear regression. Beyond head motion, compliance with PEER scan instructions (to fixate on the stimuli) can impact model accuracy. While we have no direct measure of this compliance in this sample, we did test for relationships between model accuracy and participant variables that varied considerably across participants and we hypothesized may impact compliance; in particular, age and full-scale IQ (FSIQ).

#### PEER Scan Image Preprocessing

We assessed PEER model validity after implementing global signal regression (GSR), a method to remove non-neuronal contributions to fMRI data (e.g. head motion). Although the neuroimaging community has not reached a consensus on the use and interpretation of GSR, it has been shown to remove global changes to BOLD signal caused, for instance, by respiration or head motion (Birn et al. 2006; Power et al. 2014; Kevin Murphy and Fox 2017; Kevin Murphy et al. 2009; Kevin Murphy, Birn, and Bandettini 2013; Fox et al. 2009). Given the increased likelihood of motion artifacts in the HBN dataset (Alexander et al. 2017), which includes participants at various points of maturation (ages 5-21) with typical and atypical brain development (e.g. ADHD), we implemented GSR on the masked eye orbit data from Scan1 data prior to model training. The preprocessed data were used to train the PEER model, which was applied to Scan2 (following identical preprocessing) to measure fit of the predicted time series with known calibration targets. We repeated this analysis with volume censoring (VC) instead of GSR (Power et al. 2014), using a framewise displacement threshold of 0.2mm on data from Scan1 prior to model training and estimation. Paired t-tests were used to compare the predictive accuracy between the original and preprocessed models.

#### Minimum Data Requirements

To establish minimum data requirements for accurate PEER model estimation, we systematically varied the number of calibration points used in model generation. Specifically, we randomly sampled N training points (N: 2-25) from Scan1 and used the corresponding images to train PEER (50 random samples were generated per N to estimate confidence intervals). In cases where there were fewer than 50 possible position combinations (e.g., 25 unique possibilities with 24 of 25 positions), some of the combinations were repeated within the 50 random samples selected. Consistent with our prior analyses using all calibration points, predictive validity for each PEER model was determined by comparing the predicted fixation time series from Scan2 with the known calibration locations.

### Validation

#### Identifying a Movie Based on Eye Gaze

We evaluated the ability of the PEER method to capture eye gaze sequences uniquely associated with a given movie stimulus. In order to accomplish this, we first applied each participant’s PEER1 model to their corresponding movie scans (*The Present* [TP], *Despicable Me* [DM]), thereby producing a participant-specific eye gaze sequence for each movie. Given that fMRI scans of DM contained 750 volumes while HBN scans for TP contained 250, each contiguous set of 250 volumes from DM was used in the following analyses. We then used an unpaired t-test to compare the level of correlation observed between differing participants’ time series when watching the same movie versus a different movie. To further quantify the level of discriminability between the eye gaze time series of the two movies, we trained a SVM classifier to predict which movie the individual was seeing based on a given eye gaze time series. The linear SVM classifier (C=100) was trained using half of the available participant datasets and tested on the remaining half of the participants. The results were assessed using a confusion matrix and ROC curves.

#### Out-of-Scanner Eye Tracker Measurement

For 248 of the 448 HBN participants included in this work, a second viewing of *The Present* was added outside the scanner at a later session in the study (the remaining 200 participants were either enrolled prior to addition of this repeat viewing to the protocol or had not completed the second viewing at the time of analysis). Eye tracking data were obtained during this viewing using an infrared video-based eye tracker (iView-X Red-m, SensoMotoric Instruments [SMI] GmbH) (Alexander et al. 2017), allowing us to compare PEER-derived eye fixations and those from the current gold standard. Eye tracking data were collected at the Staten Island or Manhattan site (sampling rate: 60 and 120 Hz, respectively). Similar to the design of the MRI data collection protocol, clips of *The Present* were shown at the end of the EEG and eye tracking collection protocol; thus, participants with poor eye tracker calibration or participants who were unable to complete the protocol were missing data for *The Present*. Of the 248 participants with eye-tracking data available, 43 participants with moderate to high levels of head motion in MRI data for *The Present* (defined by mean FD > 0.2mm) were excluded from the analysis. An additional 89 participants with missing eye tracking data using a stringent 10% threshold (from eye blinks or time off the screen) were removed from the analysis, leaving 116 participants in the dataset. The median fixation time series of these participants from PEER was compared to that of eye tracking, using Pearson’s r to assess the similarity between the eye gaze sequences detected by each method.

#### Identifying a Participant Based on Eye Gaze

We examined the similarity and test-retest reliability of the eye gaze time series measured by out-of-scanner eye tracking and PEER in the same participants during two different viewings of the *The Present*. To accomplish this, we calculated the intra- and inter-individual variability in viewing patterns by calculating Pearson’s r between all pairs of PEER and eye tracking fixation time series to assess the feasibility of identification. Then, we computed the correlation for each participant’s PEER-estimated and eye tracking fixation time series when compared to the median fixation time series. Using these measures, we ran a univariate ICC analysis for the whole scan and for individual time points. We also completed a multivariate extension of ICC named the Image Intra-Class Correlation Coefficient (I2C2 (Shou et al. 2013)) with 500 permutations to estimate the null distribution.

#### Neural Correlates of Eye Gaze Sequences

We tested the feasibility of using the PEER-derived fixations from movie viewing fMRI data to characterize the neural correlates associated with eye movements. Based on prior work, we expected to observe activations in the frontal eye fields, ventral intraparietal sulcus and early visual processing areas (Choi and Henderson 2015; Corbetta and Shulman 2002; Marsman et al. 2012; Bruce and Goldberg 1985); deactivations were expected in the default mode network during movie viewing (Greicius et al. 2003). To model BOLD signal activity associated with fixations identified by PEER, we first computed the magnitude of change in eye gaze location from one time point to the next in each predicted PEER time series using Euclidean distance. The resulting stimulus function was convolved with the double-gamma hemodynamic response function with added temporal derivatives, which was then used to model voxel-wise activity in response to movie viewing. To minimize the impact of prediction errors, a hyperbolic tangent squashing function was used to identify and reduce spikes in the eye movement vector prior to the convolution. Twenty-four motion parameters were regressed out from the model and FSL FEAT was used for all individual-level analyses. To assess group activation, we used the FSL FLAME mixed effect model with the following variables as nuisance covariates: sex, mean framewise displacement, age, model accuracy in the x- and y- directions and full scale IQ. Multiple comparison correction was carried out using Gaussian Random Field theory, as implemented in FSL (Z > 3.1, p < 0.05, corrected).

## RESULTS

### Validation with Simultaneous Eye Tracking and PEER

We compared the PEER model-generated eye gaze time series with that measured by a simultaneous in-scanner eye-tracker during 1) a repeat presentation of the PEER calibration task, and 2) the movie *The Present*.

For the PEER calibration scan, the correlations between the PEER model generated time series and those from simultaneous eye tracking was generally strong, ranging from .96 ± .01 (median ± median absolute deviation [MAD]) and .92 ± .02 in the x- and y- directions (Figure 1A). Comparison between eye gaze points estimated by PEER modeling and simultaneous eye tracking demonstrated high spatial concordance, with median difference (± MAD) of 1.50° ± 0.71° and 1.30° ± 0.37° in the x- and y- directions, respectively. For viewings of *The Present*, correlations between PEER-derived estimates of eye gaze location and simultaneously acquired eye tracking data were high in the x-direction (.82 ± 0.09) and moderate in the y-direction (0.59 ± 0.04). Estimates of angular deviation between the stimulus locations and PEER-derived fixations showed a median difference (± MAD) of 1.34° ± 0.34° and 1.51° ± 0.71° in the x- and y- directions, respectively. For reference, a direct comparison of PEER and eye tracking time series for one participant’s viewing of *The Present* is provided in Figure 1B, with all time series available in the Supplementary Figures 2 and 3. Individual error measures are available in Supplementary Tables 1 and 2. While angular deviation was used as our primary measure to evaluate distance, we also examined Euclidean distance, which yielded highly similar results despite being more impacted by outliers (see Supplementary Figure 4 for details).

**Figure 1.**
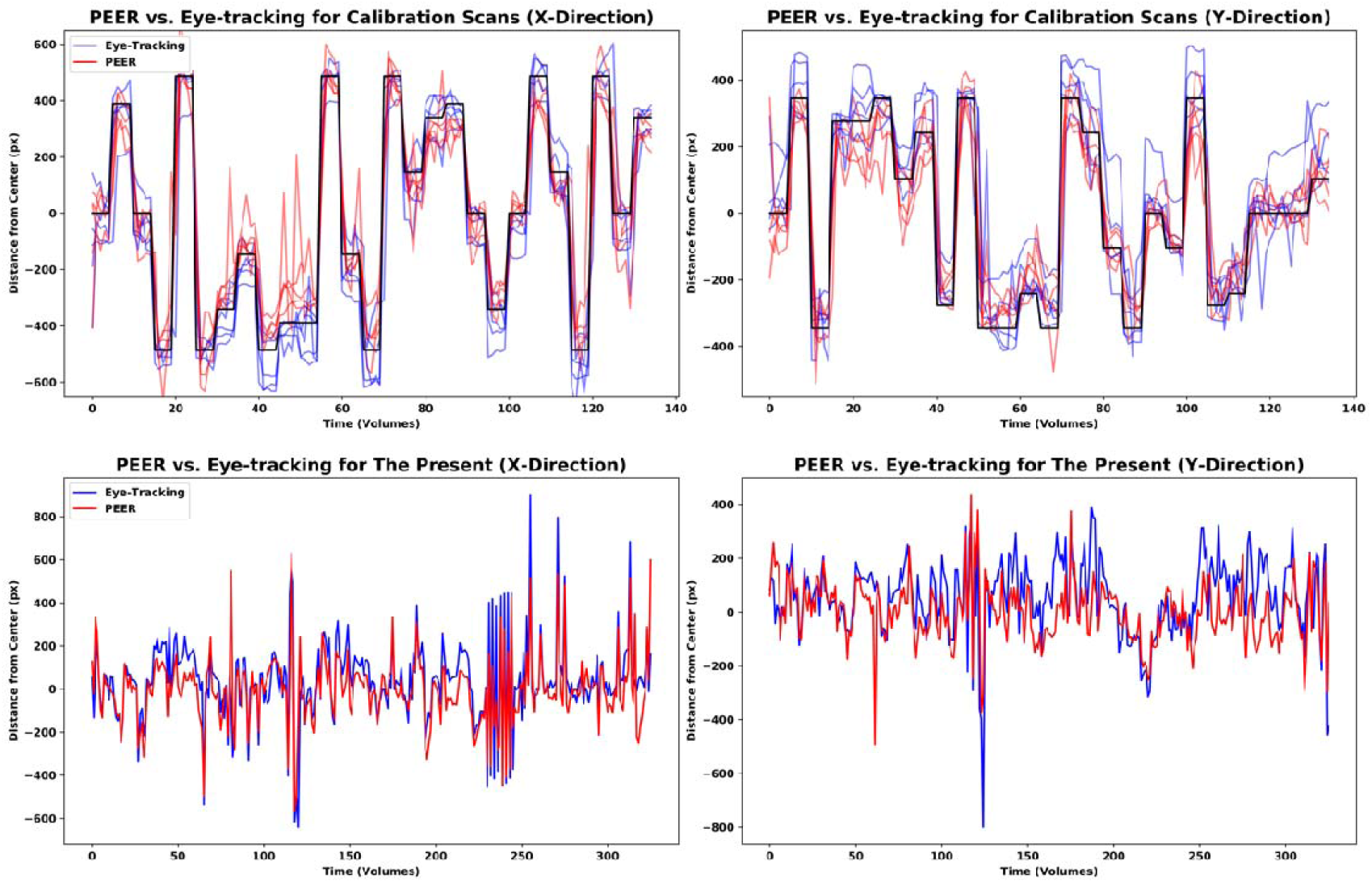
Comparison of fixation timeseries from simultaneously acquired eye-tracking and PEER predictions. (A) Here, we depict predictions in the x- and y- directions for all five participants for the PEER calibration scan (in red), simultaneous eye-tracking locations in blue, and target stimulus locations in black. (B) Eye tracking data and PEER-generated eye gaze predictions for one participant for a viewing of *The Present*. Scans for *The Present* collected at NKI included the movie credits and are thus slightly longer (330 volumes) than scans collected at HBN (250 volumes). The Pearson r values for this participant are .82 and .61 in the x- and y- directions respectively.

### Predictive Validity: Inter-individual Variation and Reproducibility

Next, we assessed inter-individual variation in predictive validity (i.e., prediction of where an individual was expected to be looking). To accomplish this, for each participant in HBN, we used the data from the first PEER calibration scan to train their SVR-based PEER model (PEER1), and then applied the model to the second PEER calibration scan to assess predictive validity. Pearson correlation scores between the predicted fixation time series with the stimulus location time series were .78 ± .21 and .68 ± .29 (median ± median absolute deviation) in the x- and y- directions (see Figure 2A for the full range of values). Assessing mismatch between predicted eye position and target locations revealed median angular deviation (± median absolute deviation across the 448 participants) in degrees of visual angle of 1.97° (±1.14°) and 1.48° (±.75°), in the x- and y- directions respectively (Figure 2B).

**Figure 2.**
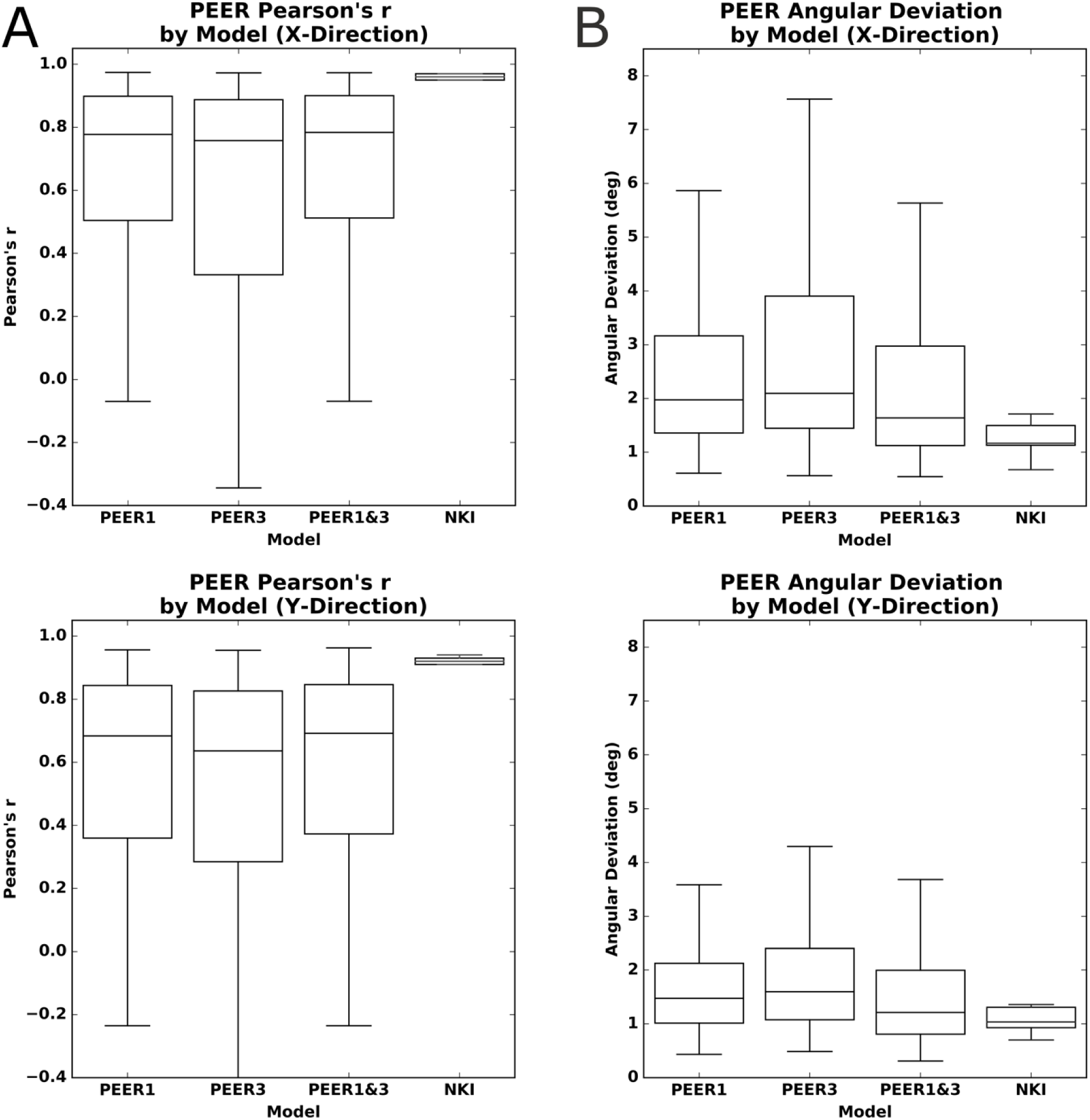
Predictive accuracy for PEER models trained on different calibration scans when estimating fixations from calibration scan 2 in the x- and y- directions. (A) Differences in predictive accuracy (Pearson’s correlation) for PEER based on the calibration scans used for SVR model training, as well as the predictive accuracy for the validation dataset from the Nathan Kline Institute [NKI]. (B) Differences in predictive accuracy (angular deviation) for PEER based on the calibration scans used for SVR model training, as well as the predictive accuracy for the validation dataset from NKI.

To assess the reproducibility of the PEER model’s predictive validity, for each participant we tested for differences when the third scan was used for training rather than the first scan (PEER3), as well as if the third scan was combined with the first scan for training (PEER1&3). We compared the predictive validities of models trained on PEER1, PEER3, and PEER1&3 calibration scans, when estimating eye fixations from the second PEER calibration scan. Paired t-tests found significantly higher validity for the PEER1 model relative to the other models (p < 0.01 in all tests), except in the y-direction when compared against PEER1&3. These differences were not dramatic (see Figure 2A) but may be explained by increased head motion, since paired t-tests showed that mean framewise displacement (FD) was significantly greater for the third PEER scan than the first (p < 0.01). We used the PEER1 model to estimate eye gaze for MRI data from the Healthy Brain Network for the remainder of the work.

### Impact of Head Motion and Age on Model Accuracy

We next considered the impact of factors that we thought may compromise model validity. First, we considered head motion, a common source of artifact in image-based analyses (Power et al. 2014; Zuo et al. 2014; Kotsoni, Byrd, and Casey 2006). We then examined age and IQ, two factors that may affect an individual’s ability to comply with task instructions. Figure 3A displays each participant’s predicted eye gaze time series for the second calibration scan plotted in rows, with the participants sorted from top to bottom in: 1) ascending order of mean FD in the training scan (i.e, Scan1) (Figure 3A left), and 2) descending order by age (Figure 3A right). Color is used to represent the predicted fixation location relative to the center of the screen and the calibration stimulus locations are depicted on top and bottom. Thus, each row represents the full predicted time series for a given participant and each column represents the predicted location for a given stimulus presentation (per TR) in the calibration scan. The consistency of fixation sequences at a given time point is made obvious by the presence of distinct vertical bands, which indicate that a highly similar fixation location is being predicted for most of the individuals.

**Figure 3.**
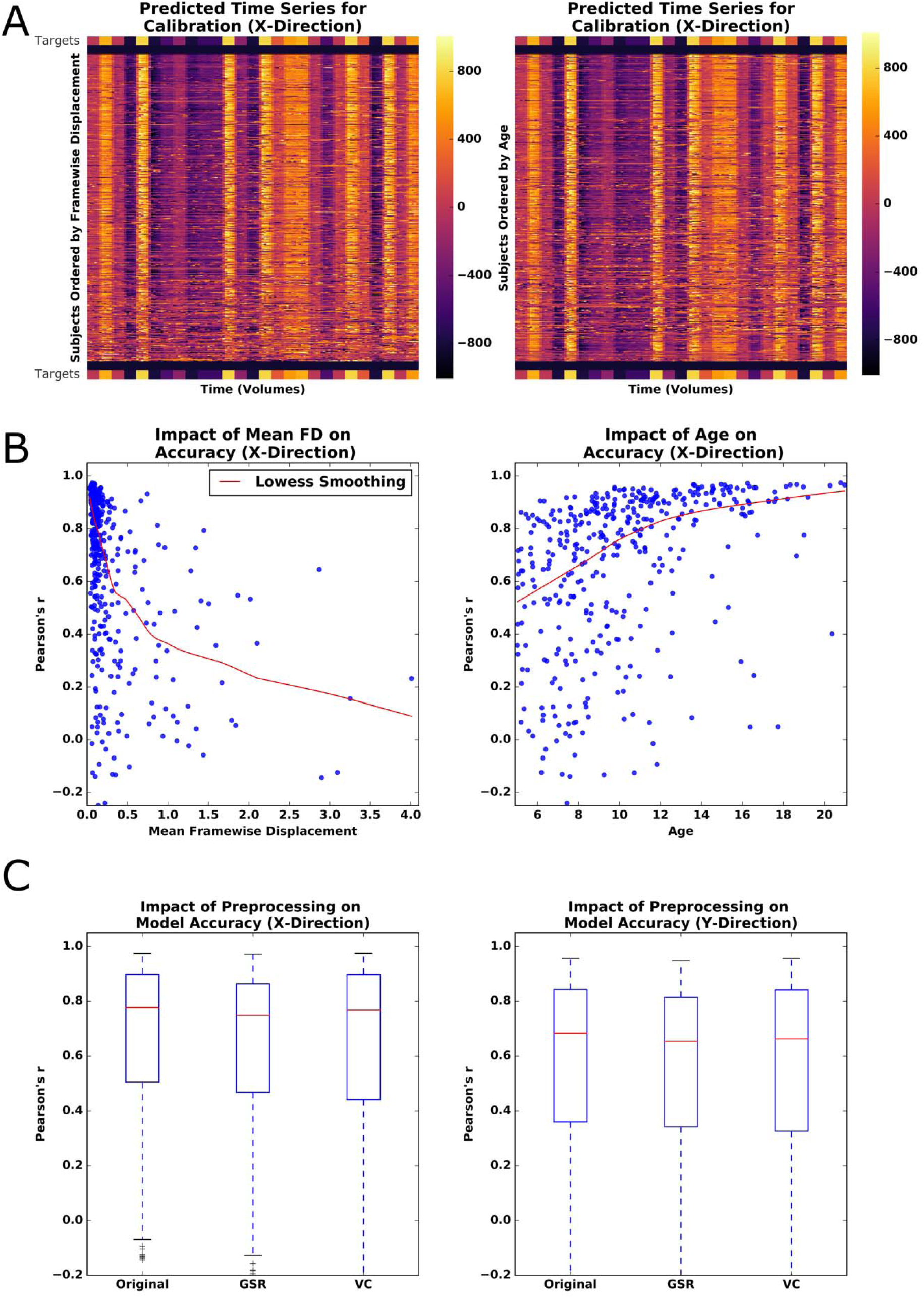
Impact of phenotype and preprocessing on PEER accuracy. (A) Heatmap of predicted fixation time series in the X-Direction from calibration scan 2, sorted by increasing motion (left) or decreasing age (right). Colorbar represents distance (pixels) of prediction from the center of the screen. Stimulus locations from the calibration scan are placed above and below participant predictions for reference. (B) Locally weighted scatterplot smoothing (Lowess) regression between PEER accuracy for the X-Direction and mean framewise displacement or age. (C) Impact of global signal regression (GSR) and volume censoring (VC) on PEER accuracy (X- and Y-Direction).

The impact of head motion (measured by mean FD) on model validity is readily discernible by visual inspection, as the eye gaze plots increasingly deviate from calibration stimulus points, moving from top (low motion) to bottom (high motion) in the plot. Findings by visual inspection were similar when sorting participants based on age, but were notably less apparent when sorting participants based on mean DVARS (D referring to the temporal derivative of timecourses, VARS meaning root mean square variance over voxels), an indirect measure of head motion, as well as other potential factors that may impact compliance (i.e., full scale IQ [FSIQ]) (Power et al. 2012). Multiple regression revealed that mean FD and age are statistically significant predictors of model validity (p < .01), but not IQ or DVARS (see Figure 3B). As expected, predictive validity is significantly negatively related with head motion (beta: x = 0.28, y=-0.27) and significantly positively related with age (beta: x=0.03, y=0.03) among those in our sample (p-values < .01 for all tests).

### Impact of Global Signal Regression and Volume Censoring on Model Validity

Given the deleterious effects of head motion, we explored whether either of two commonly used correction strategies would improve the predictive validity of PEER models: global signal regression (GSR) (applied to the orbit) and volume censoring (framewise displacement > 0.2) (Power et al. 2014; Kevin Murphy, Birn, and Bandettini 2013; Power et al. 2012). Both of these methods significantly decreased model validity when applied to the training scan according to paired t-tests (p < .01), though the differences were relatively minimal (see Figure 3C). Limiting the GSR analysis to only those participants with high motion (i.e., mean FD > 0.2) did not increase its impact on model validity; instead, no difference was found.

### Minimum Data Requirements

Previous implementations of PEER have been few in number and differed with respect to the number of target locations for training (e.g., 9 vs. 25) (LaConte et al. 2007; Sathian et al. 2011), leaving questions regarding appropriate data requirements unanswered. To empirically evaluate the impact of the number of locations included in training on PEER model predictive validity, we created random subsets of our data for each participant that systematically varied the number of training points included (i.e., from 1 to 25; 50 random subsets per number of locations). We found that PEER model validity appeared to asymptote with as few as 8 calibration targets; minor increases in validity were noted with additional training points. This is consistent with our findings of that volume censoring had minimal impact on predictive validity and that addition of a second PEER scan during training did not result in substantial increases in model performance.

### Eye gaze patterns during movie viewing

We next applied PEER to the movie fMRI scans. Specifically, the PEER1 model was used to predict fixation sequences for two scans during which video clips were viewed — *Despicable Me* and *The Present*. Figures 4A and 4B depict the predicted time series for the two movie scans. In each set of time series, distinct vertical bands were noted, indicating consistency in fixation location across participants at many of the time points, similar to what was observed for the calibration sequences. Individual timeseries are plotted (with the group median gaze pattern for reference) in Supplementary Figures 5 and 6. This is unsurprising since visually and emotionally salient scenes throughout the movie are expected to automatically capture a person’s attention (e.g. a human face in the center of the screen), while those that are less important to the narrative will be less consistent in viewing patterns across participants (Hanke et al. 2016; Dorr et al. 2010; H. X. Wang et al. 2012).

**Figure 4.**
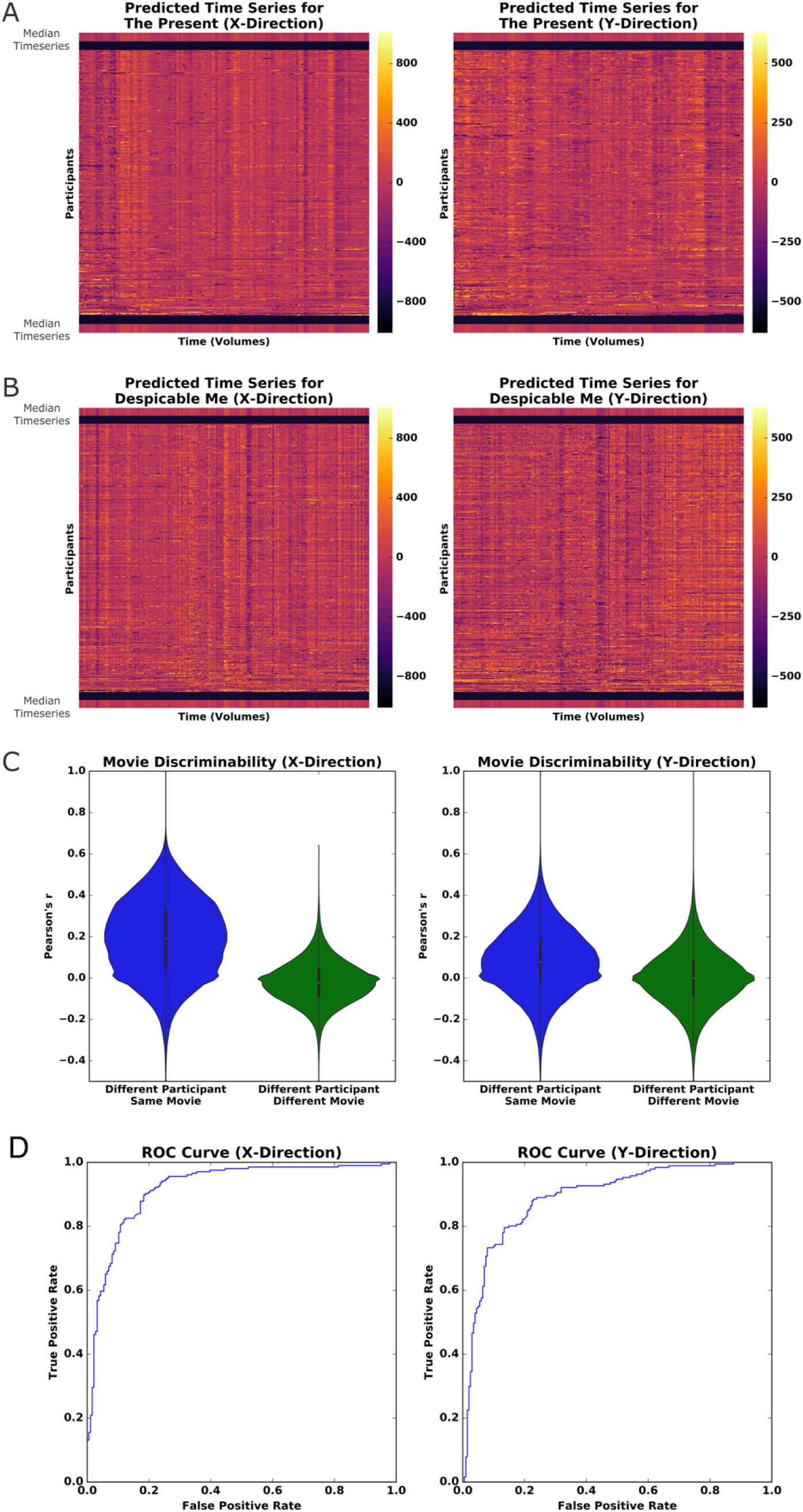
PEER applied to movie viewing fMRI. (A) Heatmap of predicted fixations for The Present. The median fixation time series is placed above and below participant predictions. (B) Heatmap of predicted fixations for Despicable Me. (C) Pairwise correlations for all predicted fixation time series from both movies sorted based on whether from Different Participant/Same Movie or Different Participant/Different Movie. (D) Receiver Operator Characteristic Curve for linear SVM model trained to identify whether a participant is watching The Present or Despicable Me based on PEER eye gaze timeseries (training carried out on half the participants; testing on the other half).

To quantify our observations, we first calculated pairwise correlations between the fixation time series generated for each of the participants for the two movies, allowing us to compare within vs. between movie relationships. As depicted in Figure 4C, correlations were significantly higher between two time series from different participants watching the same movie than correlations between different participants watching different movies. Correlation scores in the x-direction ranged from .189±.185 (mean ± standard deviation) for different participants viewing the same movie and from -.021±.103 for different participants viewing different movies. In the y-direction, scores ranged from .082±.157 for viewing the same movie and from -.002±.133 for viewing different movies. Unpaired t-testing revealed significance values < .01 in both x- and y-directions, and effect sizes (measured by Cohen’s d) of 1.14 and 0.55 in the x- and y-directions respectively. To quantify the level of discriminability between movies based on the predicted series, we trained a linear SVM using data from half of the available participants and tested on the remaining half. To match the duration of The Present (250 volumes), our analysis focused on only 250 contiguous volumes selected from Despicable Me (750 total volumes); to ensure the generalizability of our findings, we repeated this analysis on all possible contiguous subsets. The model performed well in distinguishing which of the two movies were being watched based on the PEER-derived fixations, with AUROC values of .903 ± .017 (median ± median absolute deviation) and .900 ± .025 in the x- and y- directions respectively. Assessment of the confusion matrices indicated that there was no class imbalance for samples that were incorrectly classified. We tested whether age, IQ or sex might distinguish the participants whose fixation sequences were classified accurately from those that were misclassified — however, no such associations were identified. Overall, these findings establish that eye gaze information from movie viewing is reliably encoded in the fixation time series from PEER, allowing us to predict which movie is viewed with a high degree of accuracy.

### Secondary Eye Tracker Validation

To evaluate the correspondence between PEER derived fixation patterns and eye tracking outside the scanner, we directly compared fixation sequences obtained for *The Present* with those measured using an eye tracker in the same participants on a repeat viewing of *The Present* (outside of the scanner on a different date). In Figure 5A, we again plot each participant’s predicted fixation time series in rows, with columns representing TRs, and color coded by eccentricity from fixation in X and Y dimensions. Comparing the median fixation series (calculated across participants) for each modality yielded correlation values of .85 and .81 in the x- and y- directions, respectively, supporting that PEER effectively captures relevant information about the gaze location on a TR-by-TR basis with high fidelity to traditional eye tracking approaches.

**Figure 5.**
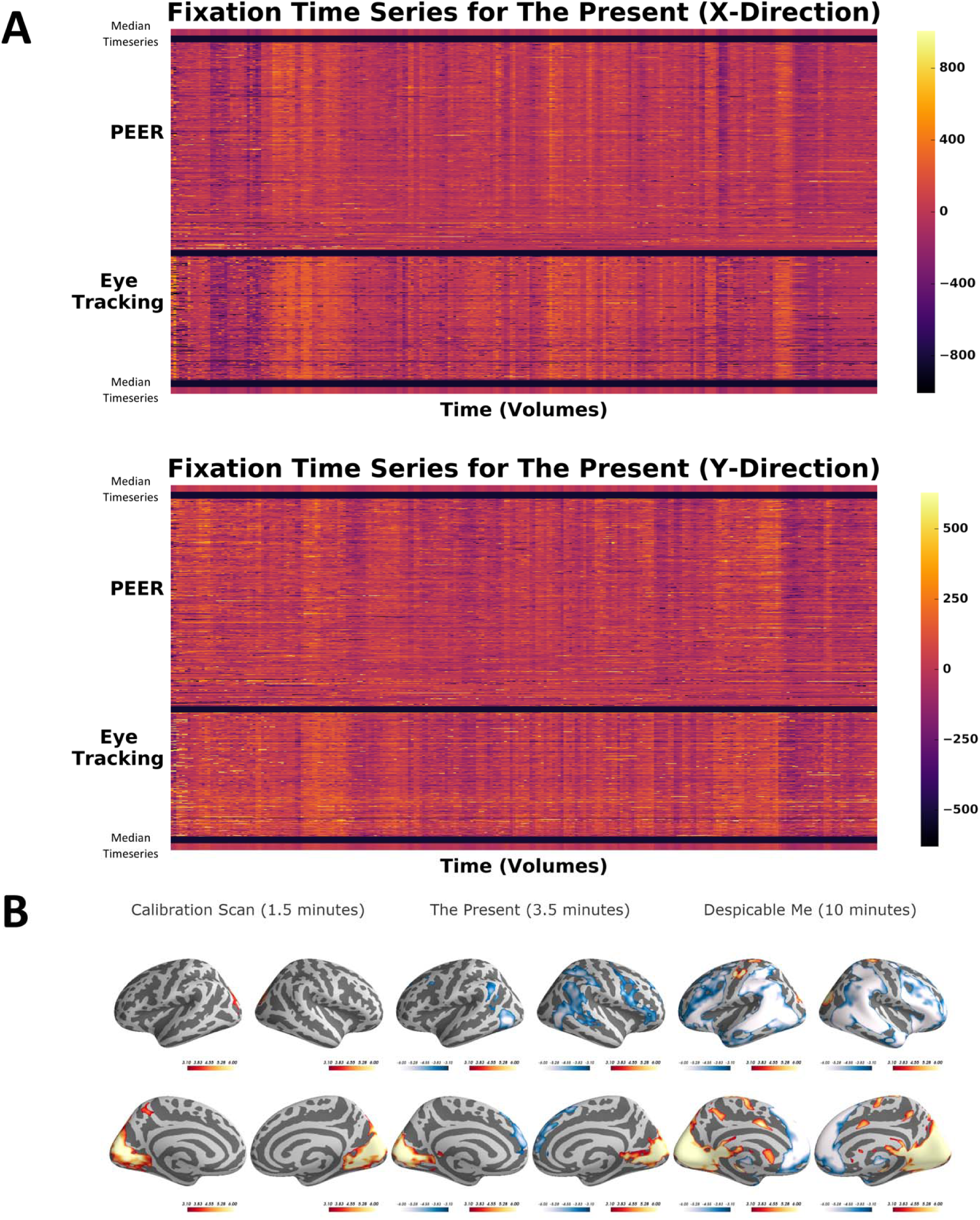
PEER and eye-tracking time series for *The Present* with activation maps. (A) Heatmap of PEER-derived fixation time series overlaid above out-of-scanner eye tracker measurements for *The Present*. The median fixation time series is placed above and below participant predictions. (B) Group activation maps generated using PEER-derived eye movements from the calibration scan, *The Present*, and *Despicable Me* (|Z| > 3.1, p < 0.05, corrected).

### Reliability of individual-specific eye gaze patterns

We first tested whether the correlation between the PEER and eye-tracker based fixation sequences were greater when collected from the same individual than different - no significant difference was detected at p < 0.05. Next, we quantified the test-reliability of fixation locations at each point in the sequence (across individuals) using intraclass correlation coefficient (ICC), as well as that of the entire fixation sequence using I2C2 (Shou et al. 2013) (a multivariate extension of ICC). Both approaches yielded poor reliability (i.e., < 0.3). The lack of reliability for individual differences may be due to the salience of the visual stimuli, which are strong enough to consistently capture attention across individuals - thereby limiting between-subject variation (H. X. Wang et al. 2012) and thus compromising test-retest reliability for individual differences. It is also possible that more precise measures derived from an eye tracker may carry meaningful individual-specific variation, though this is beyond the scope of the present work.

### Neural Correlates of Eye Movement

Finally, we examined the neural correlates of eye movements, as indexed by changes in fixation location during each of the scans (i.e., calibration, *Despicable Me, The Present*). While the low temporal resolution of PEER clearly limits the ability to identify individual saccades, or sustained smooth pursuits, we expected that it should afford a gross perspective of neural activity associated with eye movements. Each participant’s fixation sequence was first converted into a change time series by calculating the euclidean distance from one time point to the next. Convolution with the hemodynamic response was then used to identify the neural correlates of eye movements; to minimize potential head motion-related saccades, we modeled 24 motion-related parameters (Friston et al. 1996) and the PEER time series itself.

We observed overlapping patterns of activation across the three scan types, with activations being most robust for the *Despicable Me* scans, likely reflecting the longer scan duration (Figure 5B). Overlapping areas of positive activation across the three scans were most notable in the lingual gyrus, calcarine gyrus, cuneus, and superior occipital gyrus, bilaterally (see Supplementary Table 3). The two movie scans exhibited significant positive activations in the supplementary eye fields of the premotor cortex, while the calibration scan did not. Overlapping significant deactivations were found for the two movies as well, most notably in the superior medial gyrus, superior frontal gyrus, inferior parietal lobule, middle frontal gyrus, and middle temporal gyrus.

## DISCUSSION

This present work demonstrates the effectiveness of the EPI-based PEER technique to accurately predict eye gaze patterns during fMRI scans using a model generated from a simple 1.5-minute calibration scan. Eye gaze information is a prerequisite for identifying variations in compliance with task demands during functional MRI scans, particularly in naturalistic viewing paradigms where there are no observable behaviors. We found that the eye orbit contains enough information to reliably predict fixation sequences from calibration scans and video clips. In addition, the PEER method’s predictive validity increased with the number of calibration targets, but stabilized after 8 targets, suggesting that 1.5 minutes of data is more than sufficient. Unsurprisingly, head motion appeared to be the primary determinant of model validity, whether during the training scan or the scan to be predicted. Eye gaze patterns were found to be highly distinct for each movie and consistent across participants, even allowing for robust identification of the movie being viewed based on PEER alone. Both simultaneous and out-of-scanner eye tracking were used to validate the results from PEER. Consistent with prior eye tracking studies, we found that consistency of the eye tracking patterns in movie viewing (Dorr et al. 2010) observed across individuals limited the ability to reliably detect individual-specific variations in eye-fixation patterns. Finally, we found that the neural correlates of eye movement identified through PEER mirror those found in literature. Thus, PEER is a cost-efficient, low-overhead method to retrospectively determine eye gaze patterns from fMRI scans.

PEER is not intended to compete with the capabilities of modern eye trackers, most of which sample at a minimum of 60 Hz and contain additional information beyond eye fixations (e.g., pupillometry) (Palinko et al. 2010). PEER is well-suited to situations where an eye-tracker is unavailable or the quality of eye-tracker data collected is compromised. PEER is a partial solution that provides additional information regarding one of the most basic limitations present in fMRI studies. As naturalistic viewing paradigms gain popularity and find more broad usage, there will be greater demand for methods that establish the validity of findings obtained in the absence of eye fixation data. The ease with which PEER can be added to any scan protocol, requiring only 1.5 minutes of data and no additional equipment or expertise, will make it appealing for many — particularly those pursuing large-scale studies. One area where future iterations of PEER may have potential advantages over eye tracking is natural sleep imaging (e.g., infants, toddlers), as detection of eye movements is not dependent on the eyelid being open and the sampling frequency should be sufficient (there are typically 15.9 eye movements per minute) (Takahashi and Atsumi 1997).

It is worth noting that beyond PEER, several prior works have explored methods to infer eye states or eye movements from the fMRI signal. Researchers have used signal intensity variations within the segmented eye bulbs to successfully predict between open- and closed-eyes conditions (Brodoehl, Witte, and Klingner 2016). Beauchamp distinguished between a control task that did not require eye movements and an eye movement task, showing increased variance from signal derived from an eye vitreous ROI when a participant performed the eye-movement task (Beauchamp 2003). Tregellas et. al. used eye movement parameters from a local registration surrounding the eyes and optic nerve to detect whether the participant was engaged in a smooth pursuit or resting task (Tregellas et al. 2002). O’Connell and Chun predicted eye movement patterns to natural images from fixation heatmaps reconstructed directly from visually-evoked fMRI activity in early visual regions (e.g., V1) (O’Connell and Chun 2018). PEER is unique in that it is capable of providing information regarding task compliance and eye movement patterns for any fMRI scan from a session based on a minimal calibration scan, and without any dependency on patterns of neural activation.

There were two key factors that reduced the validity of the PEER model obtained. First and foremost is the deleterious impact of head motion. Consistent with observations from the resting state fMRI literature, we found higher variability in the accuracy for model estimates from data with mean framewise displacement exceeding 0.2mm. Real-time motion detection systems in fMRI could be used to help ensure the sufficiency of data quality obtained and resultant predictive validity at the time of acquisition (Dosenbach et al. 2017). Second is compliance with the calibration scan. Similar to any eye tracking paradigm, failure to comply with the calibration will compromise detection accuracy. Not surprisingly, we found that model accuracy was predicted by age (after controlling for head motion) — a finding that possibly reflects lower compliance with instructions in young children (Byars et al. 2002). One could potentially enhance and confirm compliance by adding a simple task into the calibration scan that requires a response (e.g., having a letter appear at each calibration location and requiring the participant to identify it). Alternatively, integration of real-time fMRI capabilities to the calibration scan could readily resolve the concern.

We found that despite consistency across participants, the eye gaze patterns did not exhibit test-retest reliability, whether using a multivariate or univariate ICC framework. This may in part reflect the nature of the comparison afforded by the present work, which required the test-retest comparison to be between a PEER-based measurement of eye fixations and an eye tracking-based measurement. We consider one of the primary challenges to be between-participant variation, as relatively consistent eye fixation patterns were detected across individuals, especially in dynamic scene viewing (Dorr et al. 2010). While studies that use summary statistics of viewing patterns (e.g. proportion of time fixating in a given region of interest) demonstrate higher intra-participant correlations, correlations are lower between the full fixation series for a given participant (Dorr et al. 2010; Rogers et al. 2018). This suggests that the eye fixations detected with PEER are primarily driven by salient visual stimuli. The relatively high level of engagement that tends to be associated with the selected video clips may also be a factor. Our findings should not be taken to infer that individual variation is beyond the window of examination afforded by the more sophisticated measures obtainable from current eye tracking devices.

Looking forward, there is potential to create a generalizable model for PEER that will enable researchers to retrospectively determine fixations from fMRI data, even when calibration data are not available. In addition, there are potential optimizations that can leverage multiband imaging to increase sampling rates. We demonstrate that PEER is an inexpensive, easy-to-use method to retrospectively determine eye fixations from fMRI data, a step toward opening new avenues of research and opportunities for a broader segment of the fMRI community.

## ACKNOWLEDGEMENTS

The work presented here was primarily supported by gifts to the Child Mind Institute from Phyllis Green, Randolph Cowen, and Joseph Healey. We would like to thank the Healthy Brain Network participants and their families for taking the time to be in the study, as well as their willingness to have their data shared with the scientific community. We would also like to thank the many individuals who have provided financial support to the CMI Healthy Brain Network to make the creation and sharing of this resource possible. Stephen LaConte would like to thank Drs Christopher Glielmi, Keith Heberlein, Scott Peltier, and Xiaoping Hu for helping to develop and test the PEER method. Jake Son would like to thank Youngwoo, Myungja, and Ickyoung for their love and support, and Drs. Perry, Swisher, Klein, and Milham for their patience and mentorship. No conflicts of interest declared.

## SUPPLEMENTARY FIGURES

**Supplementary Figure 1.**
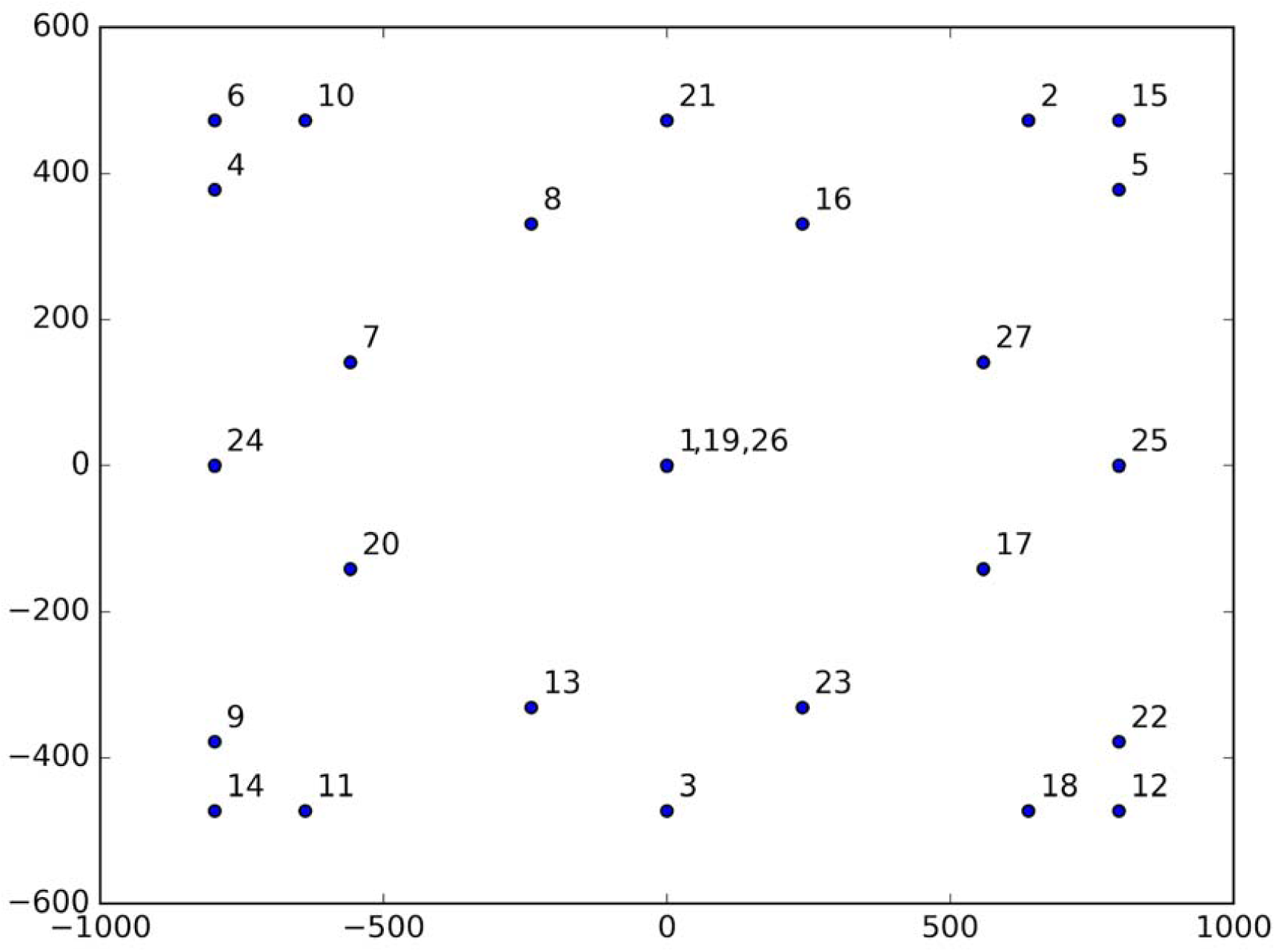
Stimulus location grid employed during the PEER calibration scans (one dot is displayed at a time in order of numbered labels; 4 seconds per dot).

**Supplementary Figure 2.**
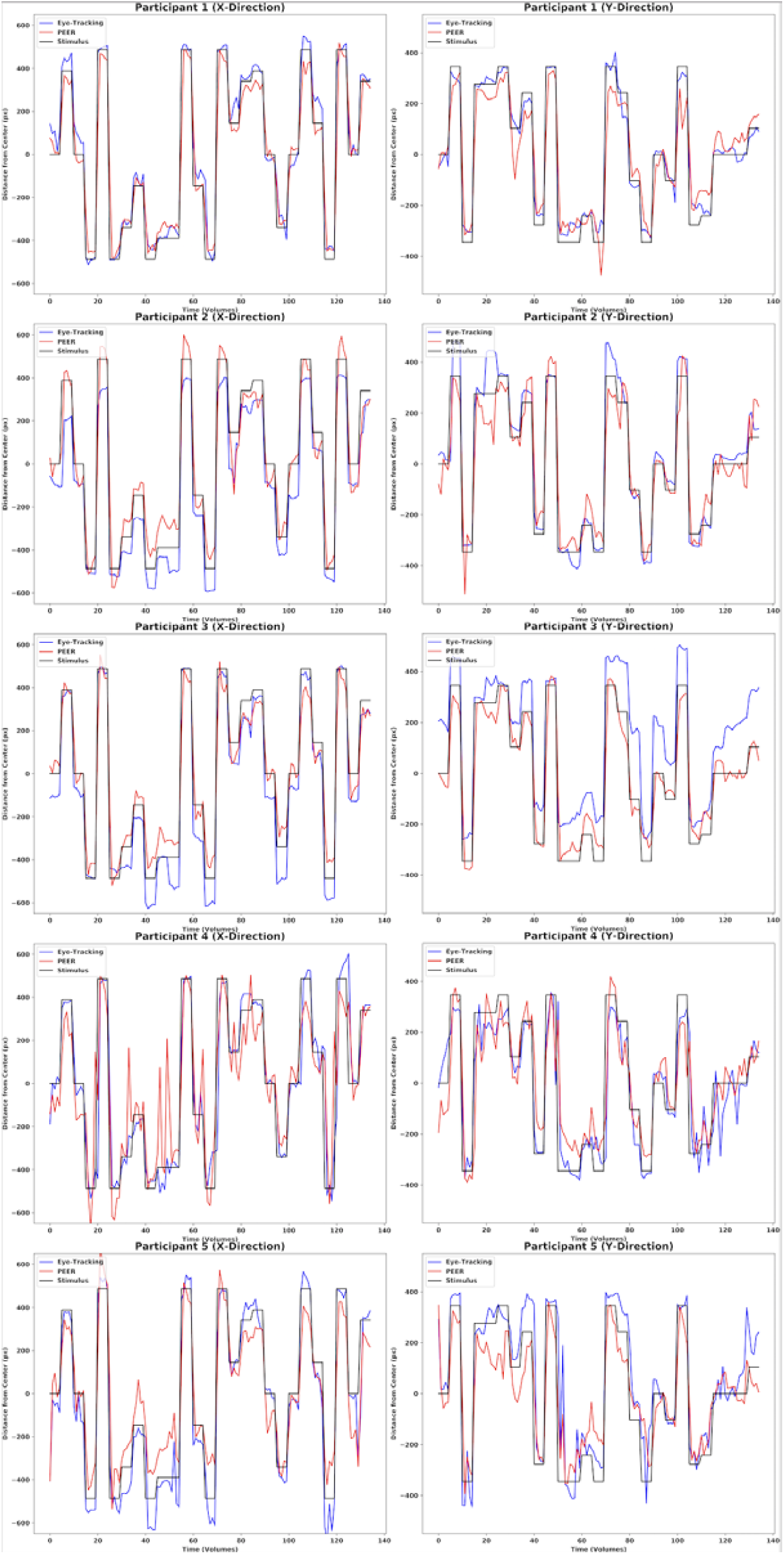
Timeseries plots of eye tracking and PEER data from a calibration scan for five participants from a simultaneously collected dataset.

**Supplementary Figure 3.**
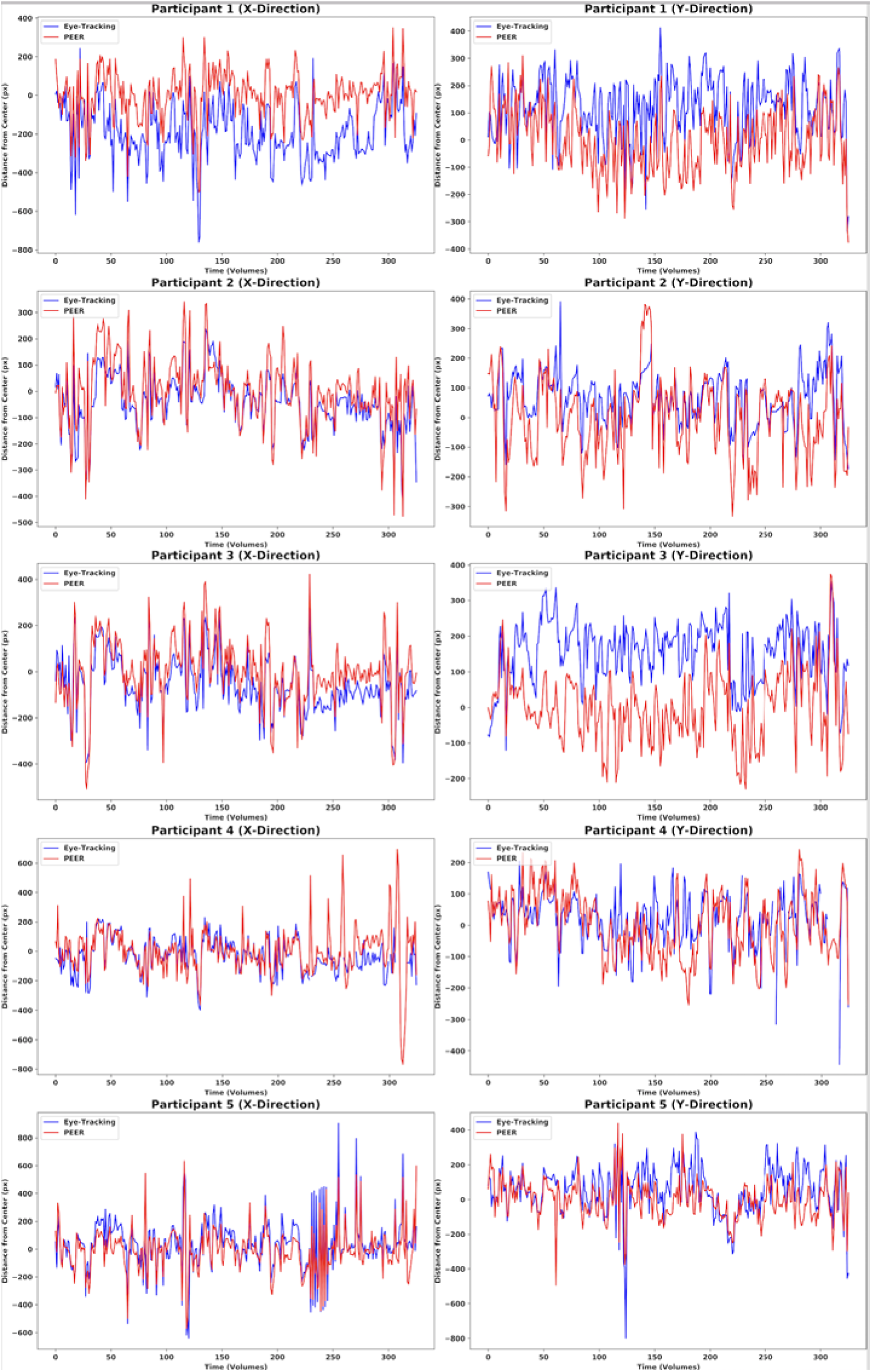
Timeseries plots of eye tracking and PEER data from viewing of The Present for five participants from a simultaneously collected dataset.

**Supplementary Figure 4.**
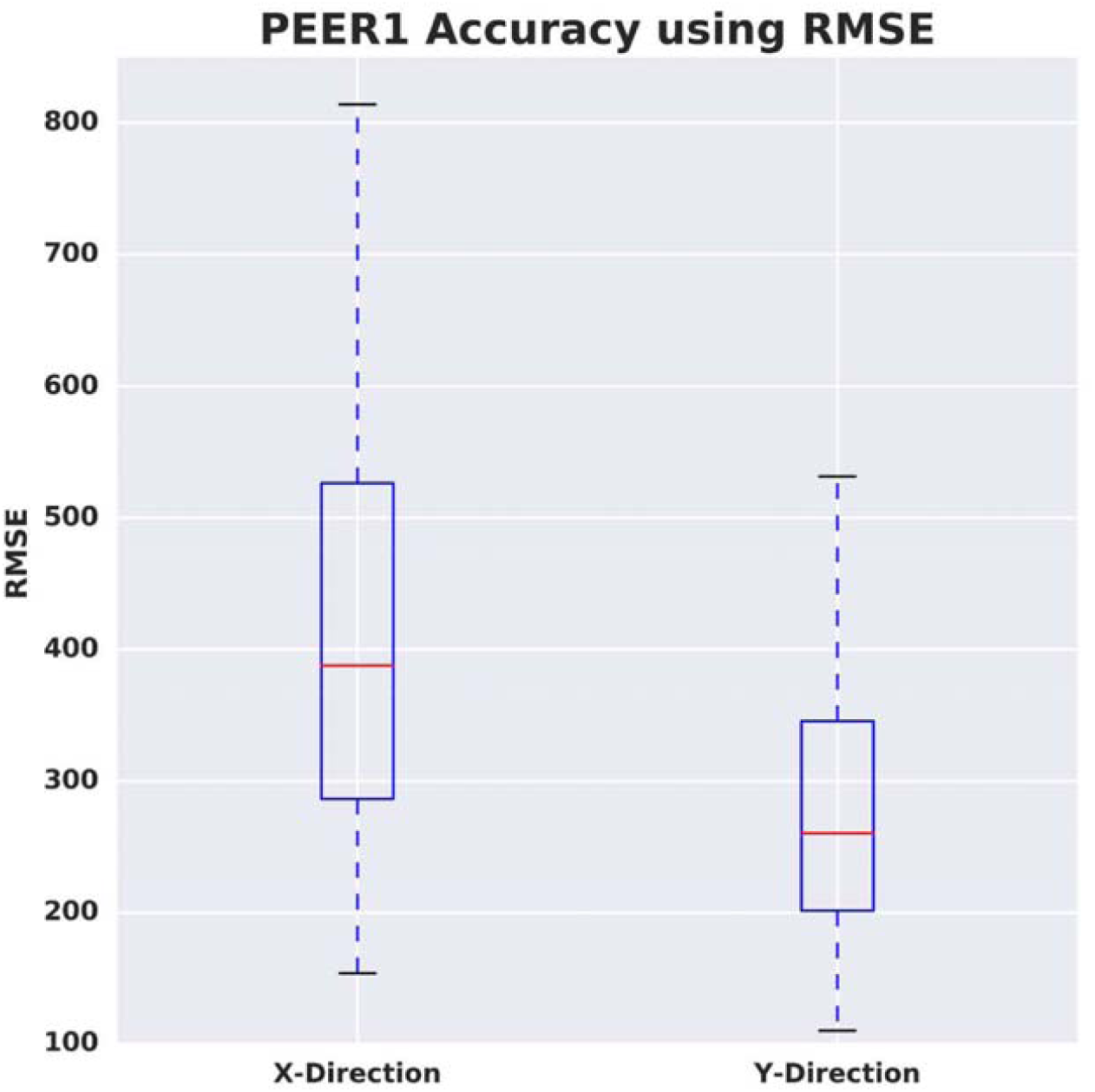
Assessment of PEER1 accuracy when estimating fixations from Scan2 using root mean square error (RMSE) of Euclidean distance.

**Supplementary Figure 5.**
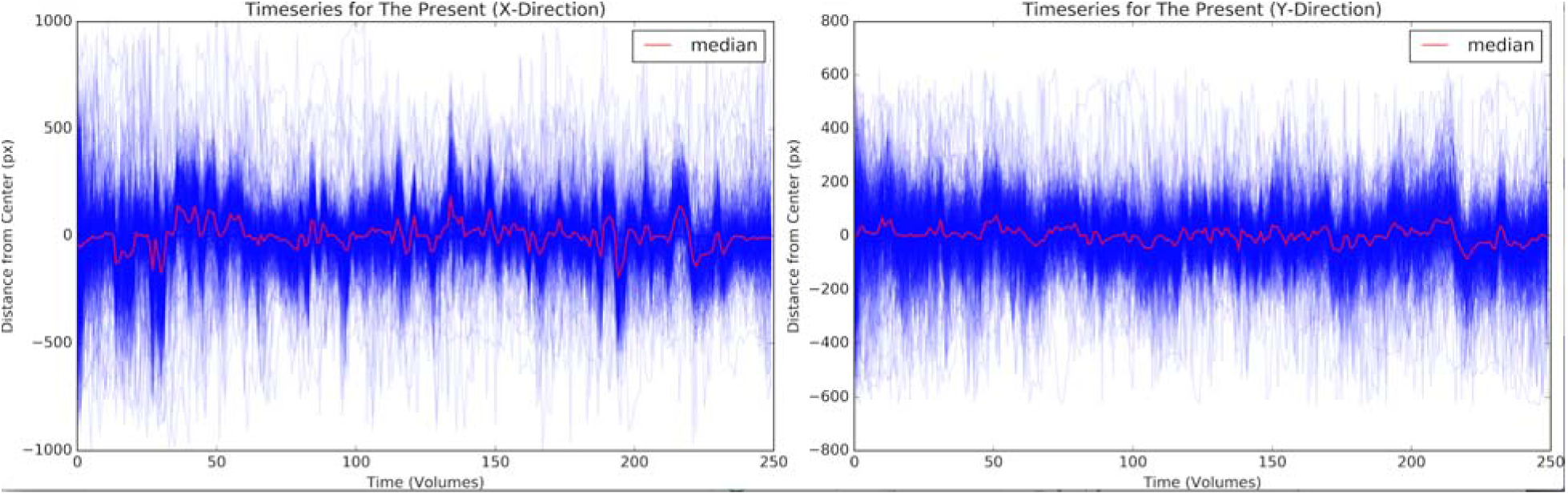
Timeseries plots of Figure 4A for *The Present*. Each blue timeseries represents a participant’s eye gaze trajectory, while the timeseries in red represents the median trajectory for the whole sample.

**Supplementary Figure 6.**
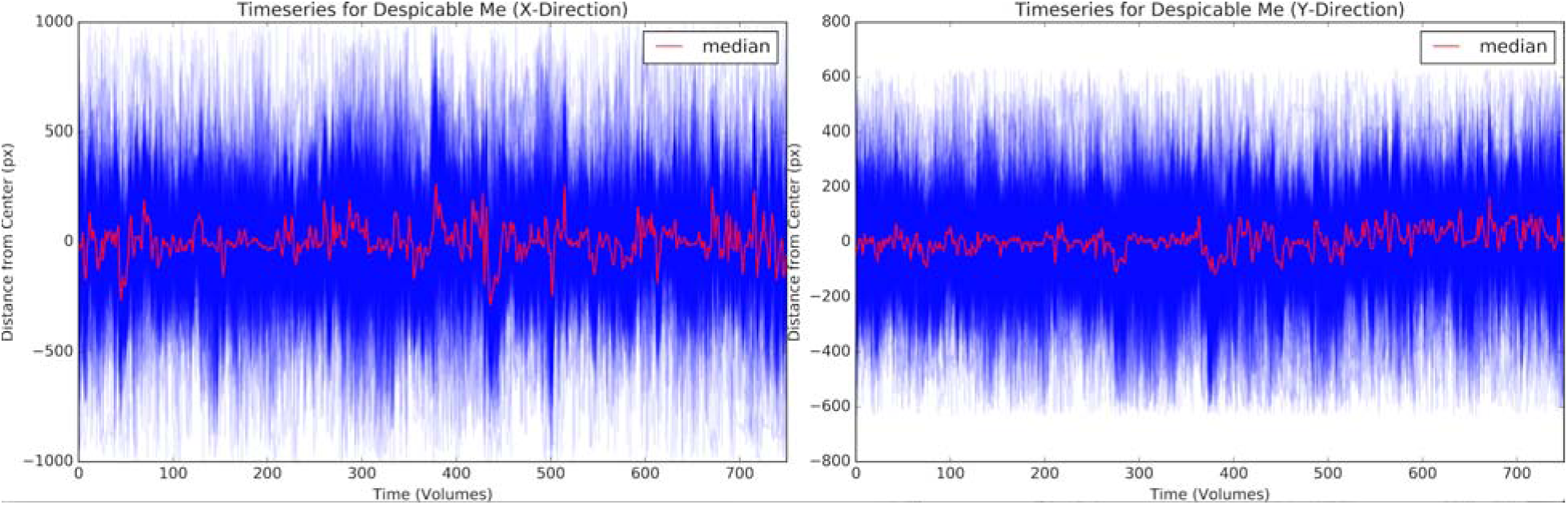
Timeseries plots of Figure 4B for *Despicable Me*. Each blue timeseries represents a participant’s eye gaze trajectory, while the timeseries in red represents the median trajectory for the whole sample.

**Supplementary Table 1.**
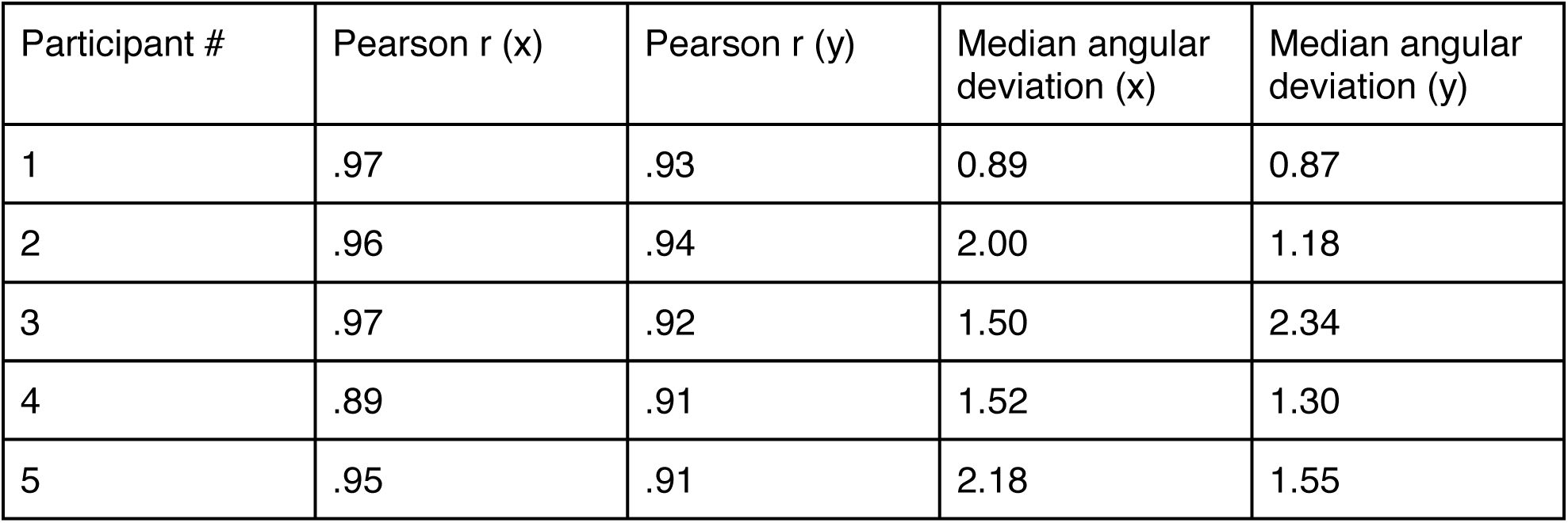
Error measures for simultaneously collected eye tracking and PEER data for the calibration scan in the x- and y- directions.

| Participant # | Pearson r (x) | Pearson r (y) | Median angular deviation (x) | Median angular deviation (y) |
| --- | --- | --- | --- | --- |
| 1 | .97 | .93 | 0.89 | 0.87 |
| 2 | .96 | .94 | 2.00 | 1.18 |
| 3 | .97 | .92 | 1.50 | 2.34 |
| 4 | .89 | .91 | 1.52 | 1.30 |
| 5 | .95 | .91 | 2.18 | 1.55 |

**Supplementary Table 2.** Error measures for simultaneously collected eye tracking and PEER data for viewings of The Present in the x- and y- directions.

| Participant # | Pearson r (x) | Pearson r (y) | Median angular deviation (x) | Median angular deviation (y) |
| --- | --- | --- | --- | --- |
| 1 | .76 | .56 | 3.99 | 2.34 |
| 2 | .87 | .66 | 0.92 | 1.51 |
| 3 | .91 | .59 | 1.34 | 3.02 |
| 4 | .66 | .54 | 0.99 | 1.03 |
| 5 | .82 | .61 | 1.22 | 1.32 |

**Supplementary Table 3.**
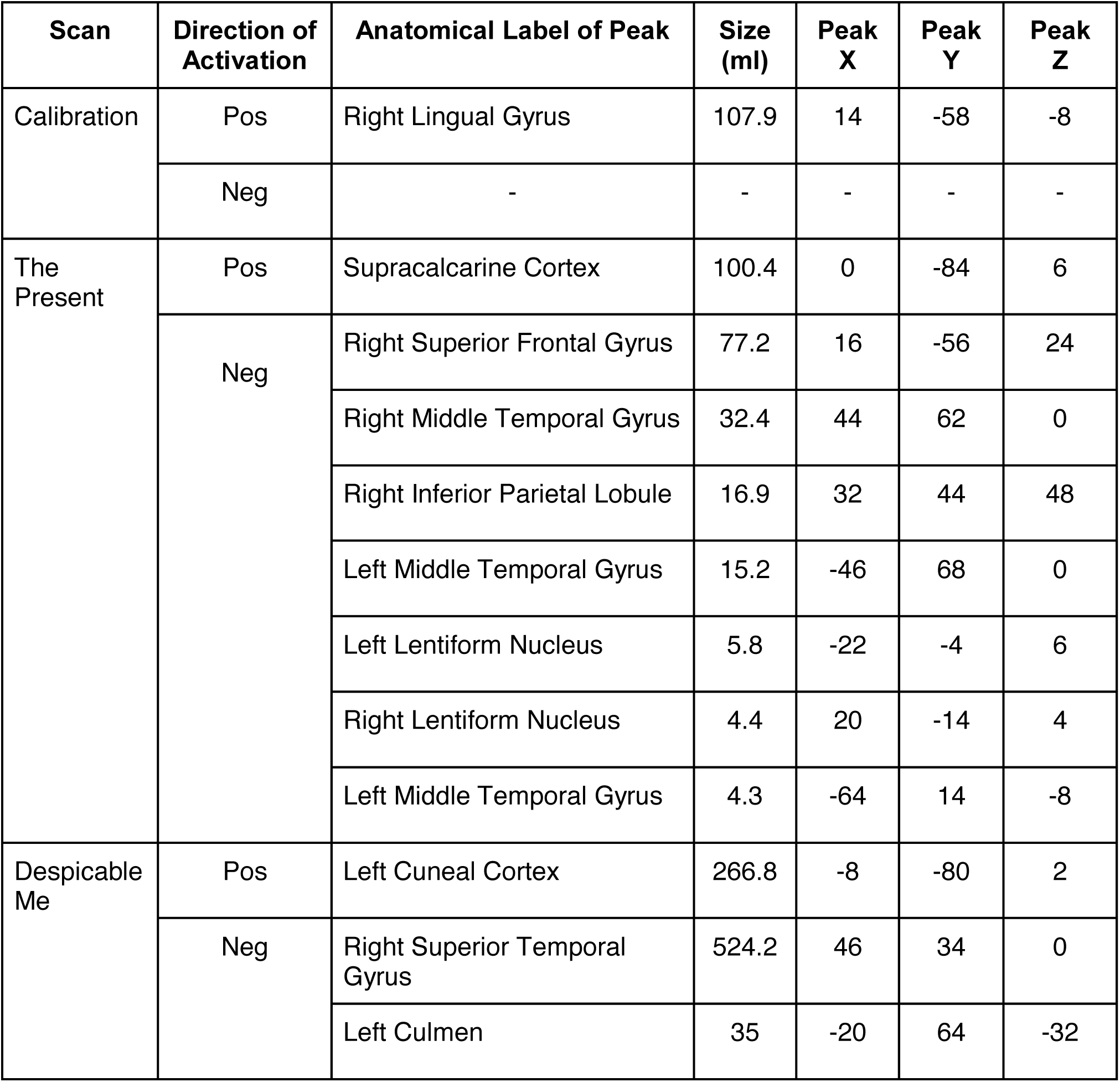
MNI coordinates of activation peaks from PEER-derived eye movements

| Scan | Direction of Activation | Anatomical Label of Peak | Size (ml) | Peak X | Peak Y | Peak Z |
| --- | --- | --- | --- | --- | --- | --- |
| Calibration | Pos | Right Lingual Gyrus | 107.9 | 14 | -58 | -8 |
|  | Neg | - | - | - | - | - |
| The Present | Pos | Supracalcarine Cortex | 100.4 | 0 | -84 | 6 |
|  | Neg | Right Superior Frontal Gyrus | 77.2 | 16 | -56 | 24 |
|  |  | Right Middle Temporal Gyrus | 32.4 | 44 | 62 | 0 |
|  |  | Right Inferior Parietal Lobule | 16.9 | 32 | 44 | 48 |
|  |  | Left Middle Temporal Gyrus | 15.2 | -46 | 68 | 0 |
|  |  | Left Lentiform Nucleus | 5.8 | -22 | -4 | 6 |
|  |  | Right Lentiform Nucleus | 4.4 | 20 | -14 | 4 |
|  |  | Left Middle Temporal Gyrus | 4.3 | -64 | 14 | -8 |
| Despicable Me | Pos | Left Cuneal Cortex | 266.8 | -8 | -80 | 2 |
|  | Neg | Right Superior Temporal Gyrus | 524.2 | 46 | 34 | 0 |
|  |  | Left Culmen | 35 | -20 | 64 | -32 |

